# Reducing error and increasing reliability of wildlife counts from citizen science surveys: counting Weddell Seals in the Ross Sea from satellite images

**DOI:** 10.1101/2020.11.18.388157

**Authors:** Leo A. Salas, Michelle LaRue, Nadav Nur, David G. Ainley, Sharon E. Stammerjohn, Jean Pennycook, Jay Rotella, John Terrill Paterson, Don Siniff, Kostas Stamatiou, Melissa Dozier, Jon Saints, Luke Barrington

## Abstract

Citizen science programs can be effective at collecting information at large temporal and spatial scales. However, sampling bias is a concern in citizen science datasets and can lead to unreliable estimates. We address this issue with a novel approach in a first-of-its-kind citizen science survey of Weddell seals for the entire coast of Antarctica. Our citizen scientists inspected very high-resolution satellite images to tag any presumptive seals hauled out on the fast ice during the pupping period. To assess and reduce the error in counts in term of bias and other factors, we ranked surveyors on how well they agreed with each other in tagging a particular feature (presumptive seal), and then ranked these features based on the ranking of surveyors placing tags on them. We assumed that features with higher rankings, as determined by “the crowd wisdom,” were likely to be seals. By comparing citizen science feature ranks with an expert’s determination, we found that non-seal features were often highly ranked. Conversely, some seals were ranked low or not tagged at all. Ranking surveyors relative to their peers was not a reliable means to filter out erroneous or missed tags; therefore, we developed an effective correction factor for both sources of error by comparing surveyors’ tags to those by the expert. Furthermore, counts may underestimate true abundance due to seals not being present on the ice when the image was taken. Based on available on-the-ground haul-out location counts in Erebus Bay, the Ross Sea, we were able to correct for the proportion of seals not detected through satellite images after accounting for year, time-of-day, location (islet vs. mainland locations), and satellite sensor effects. We show that a prospective model performed well at estimating seal abundances at appropriate spatial scales, providing a suitable methodology for continent-wide Weddell Seal population estimates.

## INTRODUCTION

Citizen science has gained in popularity as a means to collect information at temporal and spatial scales otherwise too expensive and time-consuming to study [1–3]. Simple and effective means must also exist to coordinate the survey effort and gather the data from numerous participants [4,5]. The growth of web-based informatics technologies has helped to simplify survey methodologies, and to amplify, better coordinate, and manage data from citizen scientist surveys [1,6]. With the help of citizen scientists, we completed a first-of-its-kind survey of Weddell seals (*Leptonychotes weddellii*) at pupping locations, through the inspection of high-resolution satellite images [7,8]. Here we evaluate the challenges associated with citizen science survey data and describe a new statistical approach for improving and validating population estimates. Our method can be broadly applied to correct inaccuracies and increase precision in citizen science surveys.

While citizen science has been used to assess wildlife population status and change [9,10], it is still unclear if citizen science can be used effectively to monitor a dynamic biological system to assist management plans [11]. Properly designed, a citizen science program could provide insights into the status of biological systems at the high spatial and temporal resolution needed the management of biological resources [11–14]. Statistical sampling theory requires that sampling error must be random and identically distributed throughout the sample in order to correctly estimate the precision and accuracy of a collected dataset. Yet, this assumption may be violated in citizen science datasets. The distribution of skill levels across space and time in citizen scientist surveys is generally unknown and if it is non-random, as can be expected, it can result in potentially large biases. The individual and overall survey effort in citizen scientist surveys may not be measured or controlled, resulting in another source of bias. Consequently, errors in the data extraction due to sampling bias may vary across space and time in non-random patterns [15]. If these spatial and temporal patterns correlate, positively or negatively, with the covariates hypothesized to explain them, we may draw misleading conclusions regarding covariate effects, or patterns of change in space and time [6,12,14,16].

For citizen science data to be reliable, especially for wildlife monitoring programs, we must provide assurances that the patterns elicited from these data do not reflect, or are not altered by, unidentified sampling bias. Several methodologies have been proposed to reduce potential bias in the data. Isaac et al. [12] classified these into two general approaches: one is to filter out outliers and highly imprecise records, while the other is to use the data or additional information on how data were collected to control for potential sources of bias. For example, Kelling et al. [14] show that the “semi-structured surveys” of eBird provide information on how the data were collected, as reported by the citizen scientists, permitting both the filtering of records and the use of the survey information to improve model fits. Similarly, several methods have been proposed to address differences in surveyor skill [12,17,18].

Because of the need for spatially-extensive datasets to effectively manage marine resources, citizen science is increasingly seen as a viable means for extracting meaningful information from very large datasets that in turn can help support science-driven management [19], a feat that otherwise would be impossible for even a team of researchers to achieve in a reasonable amount of time. We sought the help of citizen scientists to count Weddell seals hauled out on “fast ice” (ice fastened to the coastline permanently or temporarily) during the austral spring (October-November) [20]. These seals were mostly pregnant females or had recently given birth. A fast-ice obligate and so-called mixed-capital breeder [21,22], Weddell seal mothers nurse pups for 6-8 weeks, during which time they cease foraging and lose 30-40% of their body mass [23,24]. The high availability of energy-dense prey is necessary to achieve their subsequent recovery [25].

There is strong motivation to develop a means to monitor the pupping Weddell seal populations in order to assess how recent environment change (including changes in fast ice availability and prey abundance) might be impacting Weddell seal distributions and population numbers. In 1997 a commercial fishery was established in the northern part of the Ross Sea [26,27], exploiting reproductively-mature Antarctic toothfish (*Dissostichus mawsoni*). The Antarctic toothfish is among the most energy-rich prey of Weddell seals [28]. It is also several times larger in mass than any other fish in the Ross Sea [29]. Thus, we are concerned about a possible substantial impact of the toothfish fishery on Weddell seal populations in the Ross Sea [25]. A monitoring program of seal populations would help elucidate these potential impacts, as well as disentangle any complicating effects of climate change on their environment, in full accordance with the ecosystem-based management goals of the Commission for the Conservation of Antarctic Marine Living Resources (CCAMLR).

The southern-most Weddell seal population is located in Erebus Bay, in close proximity to the U.S. Antarctic Program’s research station, McMurdo, which permits the population’s long-term ground-based study [30,31]. Monitoring populations elsewhere would necessitate costly logistical constraints if conducted via traditional survey methods [32–35]. We devised a method to engage citizen scientists in counting Weddell seals from high-resolution satellite images of Erebus Bay and ultimately from images for the entire continent [36]. We used an online platform to enable volunteers to inspect high-resolution images of fast ice [37,38] and to “tag” (i.e., mark in the images) all presumed seals. Because the seals suckle their young while hauled out on the fast ice, they appear as “gray commas” on a white icy backdrop; thus, they are distinct enough to be detected and counted, especially as they are large enough to occupy several pixels in high-resolution images [8,33,36,37]. Moreover, while they do aggregate at predictable locations to raise their pups (henceforth “haul-out” aggregations), unlike most pinnipeds they space themselves rather than clump closely. Thus, they lend themselves to be counted using very high resolution satellite imagery. Establishing the precision and accuracy of the citizen scientist counts would allow us to determine if our approach is useful for monitoring Weddell seal populations, and at what spatial scale is it possible to then make statistical inferences about the impact, if any, of the fishery and other biophysical processes.

The work presented here has two objectives: estimating and correcting the bias and increasing precision of citizen scientists’ counts, and then calibrating the estimated counts to actual numbers on the ground. Regarding surveyor bias, our primary concern is the error associated with the possible non-random distribution of surveyor skills and effort across space, and consequent bias in estimating seal numbers. In our study of Weddell seals, the bias arises from the differential skill of taggers in correctly identifying seals in images. Some taggers are conservative and fail to tag seals in an image (omission error), while other taggers err on counting more than just seals (commission error), tagging other features such as shadows, rocks, etc. We first explore if peer ranking by itself (“collaboratively filtering”, or “crowd wisdom”) [39] is an effective means to increase the accuracy of estimates. Assuming that error due to taggers failing to tag seals is very small or nonexistent, we may filter for only the features where all or most taggers converged. Doing so we may obtain an accurate number of seals in the image, presumably filtering out odd tags on non-seal features. But even if the assumption is correct, there is no clear way to determine how the number of tags on a feature can be used to indicate it is a seal. More importantly, some non-seal features may be tagged by commonly inaccurate surveyors. To solve these two problems, a “peer-ranking” of the taggers can be used to weigh the likelihood of a tagged feature being a seal. The tagger ranking hypothetically makes tags from those taggers that are higher-ranked weigh more than tags made by taggers of lower ranking. This method allows for some features to be counted as seals even if not all taggers placed tags on them, if they were tagged by the higher-ranked taggers, and conversely, weigh down counts by lower-ranked taggers on non-seal features. We compared the above “crowd wisdom” approach to a method that corrects for potential bias using information on surveyor quality, by comparing our citizen scientist surveyors’ tags against those of an expert (author MLR) with many years of experience visiting seal haul-out locations on the ground, by aircraft (airplane and helicopter), and through inspecting aerial and satellite images for thousands of hours. This experience helps her accurately distinguish shadows and rocks from seals in the satellite images.

After reducing surveyor bias, in the second part of this study, we focused on the estimation of seal abundance as determined by direct on-the-ground surveys conducted by field biologists. We evaluated the effect of the following factors on the variance in citizen scientist counts in relation to ground counts: sensor type, time of the day, haul-out location, and seasonal period (i.e., early versus late in the pup-raising period). Since seals follow a diel pattern of foraging in which they leave the fast ice surface for short periods during the day, we expect that counts will vary with time of day [40]. Because haul-out locations differ in characteristics such as location of the cracks in relation to the sun or the sensor, and extent of non-seal features that may hide the seals from view [37], we also evaluated a site effect in the estimation of seal count accuracy. Lastly, as the nursing period advances and pups grow in size, they may become more visible in the images, so we evaluate if numbers of tags placed increase in the counts of seals in images acquired later in the pupping season [37,40]. We test for the presence of all these effects, correct for them where appropriate, and then compare the estimated, corrected number of seals to ground counts at each haul-out location and for all of Erebus Bay, and show that our approach can be used to effectively count seals at regional and continental scales. We then discuss the feasibility of using our citizen science approach for monitoring Weddell Seals throughout all of the fast ice coastal areas of Antarctica. We also discuss the general applicability of our correction method to count surveys by citizen scientists.

## METHODS

### Study area

Erebus Bay, situated between Ross Island and Victoria Land in the southwestern portion of the Ross Sea (Antarctica), is home to a > 50-year long investigation of Weddell seals, the longest-running study of any marine mammal in the world [30,31] (Fig 1). Erebus Bay is approximately 400 km^2^ and is covered by fast ice during winter that typically persists into summer months due to its sheltered embayment and southerly location [38]. Approximately 10-11 locations within Erebus Bay [37], which we aggregate into 8, represent traditional haul-out spots (used year after year) by Weddell seals for pupping. These locations primarily occur along cracks that form along the coast and extend from small rock outcrops (islets) and the Erebus Glacier Tongue. The cracks allow seals to enter and exit the water, portions of which are kept ice-free by the seals gnawing and breaking-up the ice [20]. We selected high-resolution satellite images of Erebus Bay that intersected in space with previously identified haul-out seal locations and in time with concurrent on-ice counts (e.g., those detailed in [31]). Given that approximately 5-8 ground surveys occur every year during austral spring, we used several dates in the month of November (when most seals are on ice lactating) with overlapping imagery for comparison. We used high-quality images (i.e., not overexposed, no striping) with little cloud cover (<20%) for our work [8,36].

**Fig 1.**
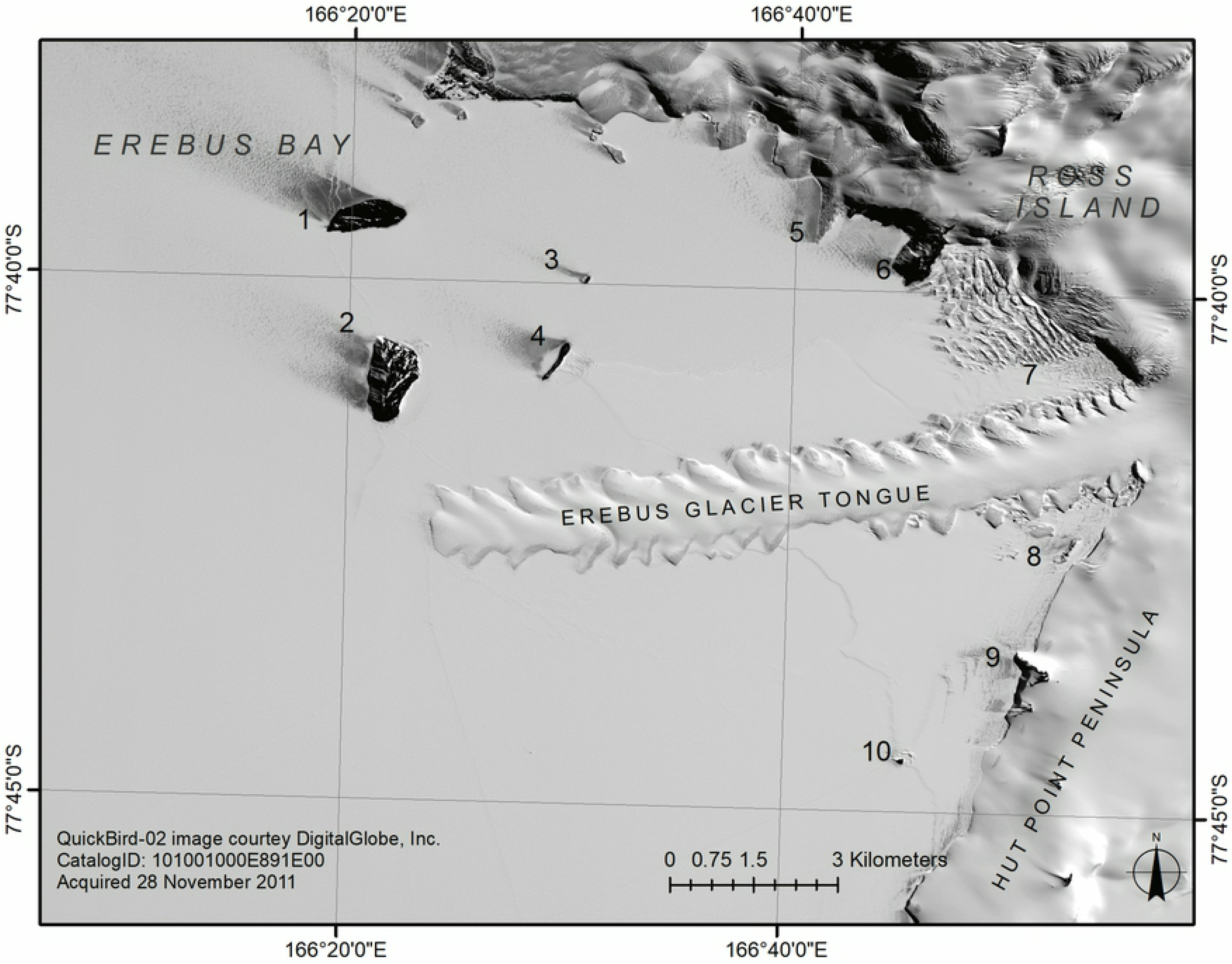
Erebus Bay study area in Antarctica. The haul-out locations (i.e., seal aggregations) studied during November of 2010 and 2011 are shown: 1. Inaccessible Island, 2. Tent Island, 3. Little Razorback Island, 4. Big Razorback Island, 5. Tryggve Point, 6. Turks Head, 7. North Base (not part of this study), 8. South Base, 9. Hutton Cliffs, 10. Turtle Rock.

### Ground counts of seals

Ground counts of seals have been conducted in Erebus Bay every year since 1969. A typical survey day includes multiple teams driving snow machines out to the seals to record the presence of all seals, to identity those individuals already outfitted with flipper tags or to affix uniquely identifiable tags to individuals not yet tagged [31]. Multiple surveys are conducted several times through a season with population estimates derived through mark-recapture methods [41]. Typically, a single survey takes approximately 12 hours to complete, and 6-8 surveys are conducted at 5-day intervals when most pups have been born and maximal numbers of adult males and females are present in the colonies during November and December.

### The citizen scientist platform

We used the Tomnod crowd-sourcing platform (now called “GeoHive”, DigitalGlobe, Inc.), which allows for efficient and effective search of large swaths of Earth’s surface [8,36,42]. Citizen scientists interested in assisting in searches logged into the platform and were each provided a unique identifying number. Instructions and training materials were provided, including multiple examples of what seals look like at multiple scales (close-up images, photographs from aerial surveys and satellite images), and hints as to where seals are likely to be found (i.e., near cracks in the ice), and other items on the landscape that could be confused with seals (e.g., rocks, melt pools). After reviewing these, participants were shown part of a satellite image on the interactive platform and were allowed to search for seals. The imaged sections shown were 500-m x 500-m in area, ~0.45-0.6 m in resolution, and included only coastal fast ice areas. Henceforth we refer to these sections as “maps.” A surveyor inspects and tags as many features s/he considers are seals in each map, and then navigates to another map that the platform selects at random from the pool of maps available for inspection. Additional description of the platform is provided in [36].

### Estimating feature and surveyor ranks, end of tagging campaigns

Through peer-to-peer ranking, the probability of a surveyor finding a “feature” (a presumptive seal) in a map is determined through comparison with data from his/her peers (an approach similar to that of [43]). We call this probability the “surveyor CrowdRank.” Formally, the surveyor CrowdRank is the probability that the surveyor will identify a feature on a map as being a seal that at least one other surveyor has also identified as a seal (i.e., the probability of tagging a feature that others have also tagged). A surveyor’s CrowdRank increases the more features s/he tags that others have also tagged. Conversely, a surveyor’s CrowdRank is reduced if s/he identified features that others did not. (There is no penalty for missing features.) Notably, the surveyor’s CrowdRank is not the probability of the surveyor identifying a seal. Since we asked that the surveyors find and tag seals, the features are presumed to be seals. We assumed that surveyors can err both in the detection of “features” (as defined above), and in the correct identification of these features as seals.

Similarly, once at least two surveyors have placed a tag on a feature, the probability of the feature *j* being detected (the “feature CrowdRank”) is calculated as from all surveyors (*i*) that tagged it, under the assumption that the surveyors made their determination independent of each other, as:

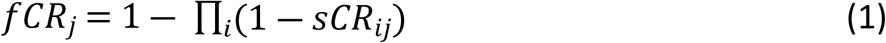

In equation 1, *fCR* and *sCR* are the feature and surveyor CrowdRanks, respectively. (A surveyor is never exposed to other surveyors’ tags previously placed on a map.) Therefore, only features with tags from at least two different surveyors are considered in the counts – that is, a feature only tagged once is not included. The estimation of the feature CrowdRank does not account for “negative” votes; i.e., surveyor CrowdRanks of those who inspected the map and did not tag the feature. Since the surveyors are looking for seals, the feature CrowdRank is really the probability that the crowd of surveyors will agree on a feature being a seal. It is intuitive to see that if two surveyors have high CrowdRank scores and both place tags on the same feature, there is a high likelihood that the feature will be deemed to be a seal by the algorithm. The surveyor CrowdRank scores are also used to determine if maps should remain in the pool to be inspected and, consequently, if the tagging campaign has been completed. The Supporting Information S1 document provides details on how the CrowdRank values are calculated and how maps are removed from the pool of available maps for inspection.

The determination of the surveyors’ agreement on a feature is done by evaluating the distance between the tags they placed every time the CrowdRanks are calculated. Because adult female Weddell seals are ~3 m long, we deemed that two surveyors agree on a feature if their tags are placed within 3 m of each other. Therefore, to set the number of features identified by the surveyors on a map, the platform identifies the maximum number of clusters of 3 m in diameter among all tags placed on the map, with the proviso that a cluster can only contain one tag per surveyor.

### Evaluating the feature CrowdRank score to reduce counting bias

Surveyors generally over-estimated the number of seals on maps [36], and often also missed placing tags on seals. The first approach to correcting the bias in counts of seals, which we evaluate here, is to apply a threshold to the set of features in a map based on their feature CrowdRank (the “wisdom of the crowd” method). We varied the feature CrowdRank threshold from 0.6 to 0.95 in increments of 0.05, obtaining a single value of the crowd’s count of seals for each map for each threshold. After filtering at each threshold level, we fit a linear model of the number of features identified by the crowd versus the expert’s counts in 601 maps (i.e., the total number of maps the expert inspected). To determine the optimal threshold, we evaluated the magnitude of the slope parameter (i.e., change in the “number of features” for the crowd versus the change in the number for the expert) and the adjusted R-square value of the regression, in addition to a visual inspection of the model fit. Since the feature CrowdRank is the probability that a feature will be deemed a seal by the crowd, we expected that both the accuracy and precision of the slope of the regression increase as the threshold used for features increases. That is, the slope of the regression approaches 1 and its standard error drops as the threshold value increases. An alternative outcome is that increasing the threshold filter would exclude an increasing number of possible features, shrinking the crowd count toward only the features with the highest probabilities of being identified as seals by the crowd. A too high threshold incurs the risk of failure to count seals that are present, which would represent a decrease in accuracy (coefficient less than 1). Our goal was to find a compromise between the accuracy in the number of seals detected by the crowd and the precision of the estimate through the selection of an adequate threshold for the features’ CrowdRank.

### A novel approach to correct bias in citizen science counts

As an alternative to just filtering out features, we explored estimating the number of seals from the citizen science counts using information both about the surveyor’s CrowdRank and how well the surveryor’s estimate compared to the expert’s (the “expert correction” method). For each surveyor, we counted the number of true and false positive tags versus actual seals, and the number of “false negative tags” (i.e., the seals that the surveyor failed to identify) in maps also inspected by the expert, on the assumption that the expert provides the true number of seals in each map. Specifically for each surveyor, given that we start with the knowledge of the correct number of seals in map *m* and number of tags the surveyor placed in that map, we can determine the individual surveyor’s probability that a tag placed is a seal (Prob[S|F] in equation 2), and the probability of not tagging a seal (Prob[!F|S]). With these probabilities, we can fine-tune, or ‘shrink’ (as defined below, see also [44]) the surveyor’s estimate of number of seals in each map s/he inspected as follows:

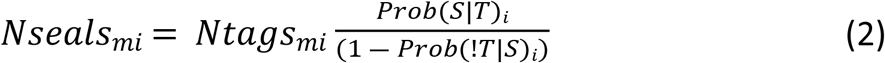

That is, the number of seals in map *m* according to a specific surveyor *i* is the number of tags s/he placed in the map, times the probability of a tag placed is indeed a seal, times the inverse of one minus the probability of there being seals s/he did not identify and tag. The first two components in the RHS of equation (2) are readily understood. The product of the number of tags in the map times the probability of a tag being a seal is equivalent to calculating the expected number of seals the surveyor tagged in a map. The denominator in equation 2 is simply an inflation factor to correct for the probability of not placing a tag when a seal is present. A detailed explanation of how we developed equation 2 and calculated the probabilities is provided in the Supporting Information S1 document.

We can then define the ratio of probabilities in equation 2 as the “shrinkage factor” for surveyor *i*. Unlike the previous “wisdom of the crowd” approach, with this method we obtain a shrinkage factor for each surveyor. This permits the calculation of a mean or median shrinkage factor from the sample of surveyors who inspected maps in common with the expert, and a measure of its precision too (e.g., the standard deviation or inner 95th quantile of the empirical distribution of individual shrinkage factors). To do this, we assume that the set of surveyors for which we can calculate a shrinkage factor is a representative sample of all surveyors involved in the campaign. This is justified since we found no evidence of any bias in the selection of surveyors.

Since the surveyor CrowdRank measures how well a surveyor agrees with her/his peers, it can act as an index of representativity among surveyors. Those with higher CrowdRank score agree more often with their peers. We again hypothesized that including only the higher ranked surveyors increases the precision and accuracy of estimates by filtering out less representative (outlier) surveyors. Thus, we examined the need and adequacy of filtering surveyors based on their CrowdRank score before estimating the sample shrinkage factor. We varied the threshold for surveyor CrowdRank scores from 0.6 to 0.95 in increments of 0.05. As with the previous filtering approach, at each threshold level we obtained a corrected estimate (i.e., after shrinkage) of the number of seals in each map and regressed it versus the expert counts. We again evaluated the adjusted R-square value of the regression along with a visual inspection of the model fit. Increasing the threshold of surveyor CrowdRank scores should result in a more precise estimate of the sample shrinkage factor, but it also means using fewer surveyors to estimate it. Doing so translates into a smaller and thus potentially less representative sample of all surveyors’ shrinkage factors, which could result in loss of accuracy. Thus we sought a threshold value that resulted in a desirable compromise between representativity (i.e., including the largest number of surveyors possible) and precision of the sample shrinkage factor.

### Effect of covariates on accuracy of counts

The two methods (“wisdom of the crowd” and “expert correction”) were used to estimate the number of seals in the maps, and the resulting estimates were then compared to the counts by our expert for the same maps. The expert correction method resulted in the best approach (as demonstrated below), and is the only one we used in the second part of this study. In order to estimate the actual number of seals at haul-out locations, once we found a suitable surveyor CrowdRank threshold to be used to shrink the counts, we obtained the necessary shrinkage factor by taking the median of the sample. We used the median because of possible outlier effects that could skew the mean value due to our low sample size.

We used the shrinkage factor to correct the sum of counts from the set of maps that encompass every known haul-out location in the study area to obtain estimates for each date surveyed. Different image dates-times were used to sum the estimates from all maps for all locations and date-times, and these sums were considered replicates of the estimated count of seals for a location if they were taken within 3 days of the ground count (see Results for details). We then ran the regression of our estimates versus the respective ground counts. The regression used the ratio of the crowd-based shrunken estimate to the actual ground count (the “estimated detection rate”) as the response variable.

We include the effects of sensor type (images come from three satellite sensors: QuickBird-2, WorldView-1, and WorldView-2), hour of the day (images were collected at various times, between 8:00 and 18:00 local time), time of month (categorized as early versus late November), year (images are from years 2010 and 2011), and haul-out location [36]. Initial regression analysis (not shown) indicated possible evidence of a difference between sensors in the detection rate. The distribution of values of detection rate is very similar between images taken by the two WorldView sensor types, but notably different for the QuickBird-2 sensor (results not shown, standard deviation of detection rate values is 0.09, 0.12 and 0.19 for WorldView-1, WorldView-2 and QuickBird-2 sensors, respectively). Thus, all counts from QuickBird-2 images were removed from the data for further analyses.

Weddell seals show a daily diving pattern foraging for brief periods while lactating [40,45]. Since daylight lasts 24 hours during the spring-summer, and most of the images were taken between 8:00 and 18:00 local time, we fitted a sinusoidal function to the hour values in local (Western Ross Sea) time. We tested several moduli (from 7 to 12 hours) for the sinusoidal pattern’s periodicity with the regression model and found modulus 12 hours to be the best fit (that is, one complete sine wave occurs over a 12-hr period). We therefore evaluated all the effects with the sinusoidal with a 12-hour modulus. Thus, the covariate used in the regression model is the value of the function:

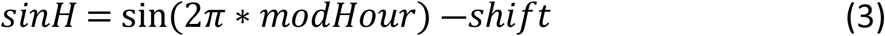

In equation 3, modHour is the hour of image acquisition in local time relative to the 12-hour modulus, and the shift parameter is the displacement of the wave (in time) to match the peak of seal presence on ice with the peak of the function. We considered two parameters that may add further information to explain the variance in the response variable: the total number of tags placed by all surveyors who met threshold and the number of maps of a haul-out aggregation where tags were placed. We expect that the more tags placed and the more maps inspected around a location, the more engaged the surveyors may be in detecting and tagging the seals. We include these as potential explanatory variables, along with their interaction, after scaling them (i.e., to a mean of 0 and SD = 1). We evaluate the significance of each of these effects on the estimated detection rate.

### Estimating the number of seals

Here we sought to develop a model that could be used to estimate the number of seals elsewhere around Antarctica, for any location where a crowd count – but no concurrent ground count – was available. To this end, we fit two final regression models to the data. One includes the effect of individual location (as a factor). A second model was fit without the effect of individual location but with a two-level factor “islet versus mainland”, which was of interest because such a model can be applied to new locations outside of Erebus Bay. Though none of the Erebus Bay locations were associated with the mainland coast of Antarctica proper, we considered haul-out locations directly on the coast of Ross Island as behaving like a mainland shore. Ross Island is many orders of magnitude greater than the small rock outcrops, or islets, where most haul-out aggregations are found in Erebus Bay. Of the 8 locations in our sample, Hutton Cliffs and Turks Head-Tryggve were along the coast of Ross Island (i.e., “mainland”), while all others were associated with peripheral islets. We distinguish these two models hereafter as the “haul-out location” and “islet/mainland” models.

The models predict the ratio of seals counted by the crowd to seals counted on the ground – i.e., the detection rate, after accounting for all relevant effects, using the expert correction method. Thus, to obtain the predicted number of seals, we divide the crowd count by the predicted detection rate:

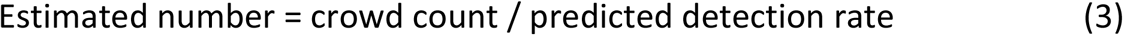

To further reduce the dispersion in the data, we fitted a Gamma distribution to the values of the detection rate under each model separately. The Gamma is a two-parameter distribution bounded at 0 on the lower end, which matches the expectation of values of the detection rate. Visual inspection of goodness of fit (provided in the Supporting Information S1 document) confirmed an adequate fit. We used values within the inner 95th quantile limits of the distribution to calculate the corrected seal numbers. We estimate the number of seals for the locations by predicting with both regression models and provide the inner 95% intervals from the standard error of model predictions. Predicting with the model with islet/mainland effect is straightforward using equation 4. To predict with the model with location, if the location is an islet, we use equation 4 and the ratio of each location that is an islet in our sample, and then average the six estimates. We do the same *mutatis mutandis* for the two locations on mainland shores. We used the average standard error of predicted values of maps within a location to calculate the confidence interval of the prediction.

Finally, we explored an approach to estimate abundances region-wide. That is, we use the islet/mainland model to predict at each location, but add up the predictions to provide a single estimate for Erebus Bay as a whole for each date-time an image was taken. We summed up estimates within each satellite image and compared to the sum of ground counts for the same area. Because the coverage of each image varies and a single image does not include the entire region, the numbers estimated comprise only the sum of haul-out locations present in the image, and the estimates are compared to the appropriate haul-out location of ground counts.

Data and analysis code showing all the analyses, tables, and graphs included in this paper are available at this URL: XXXXXXXXXXXXXXXXXX

## RESULTS

Overall, 241,478 surveyors inspected 292,049 maps from Erebus Bay and placed 128,061 tags in them. Since these images were taken during 53 different times on two different years, each survey of Erebus Bay required the crowd to inspect on average 5,510 maps, or the equivalent of 1,377.5 km^2^. Most maps had no seals or features on them. Indeed, tags were placed only on 12,937 maps (4.4% of all maps inspected), with nearly 2/3 of these having only single tags. The expert (MLR) also inspected 10,143 maps from a subset of 23 images. The expert found a total of 2,862 seals, but in only 525 (5.2%) of maps inspected. Hence, the ratio of maps with tags versus total inspected between the taggers and the expert was comparable (4.4% versus 5.2%).

In an initial campaign [36], we asked surveyors only to indicate if a map contained seals or not (yes/no answers, not counts). In this case they only missed 3 of ~20,000 maps where seals were present (as confirmed by the expert). In all subsequent campaigns, surveyors were asked to count each seal (i.e., place tags on each seal found). However, surveyors commonly placed more tags than the number of seals present in a map. This is readily shown in Fig 2, which plots the expert count versus the crowd estimate of “features” (i.e., those that received at least 2 tags). Most crowd estimates are higher than the expert count (commission error). Also, though they very rarely missed identifying maps with seals, individual surveyors were likely to miss tagging some seals in these maps (omission error).

**Fig 2.**
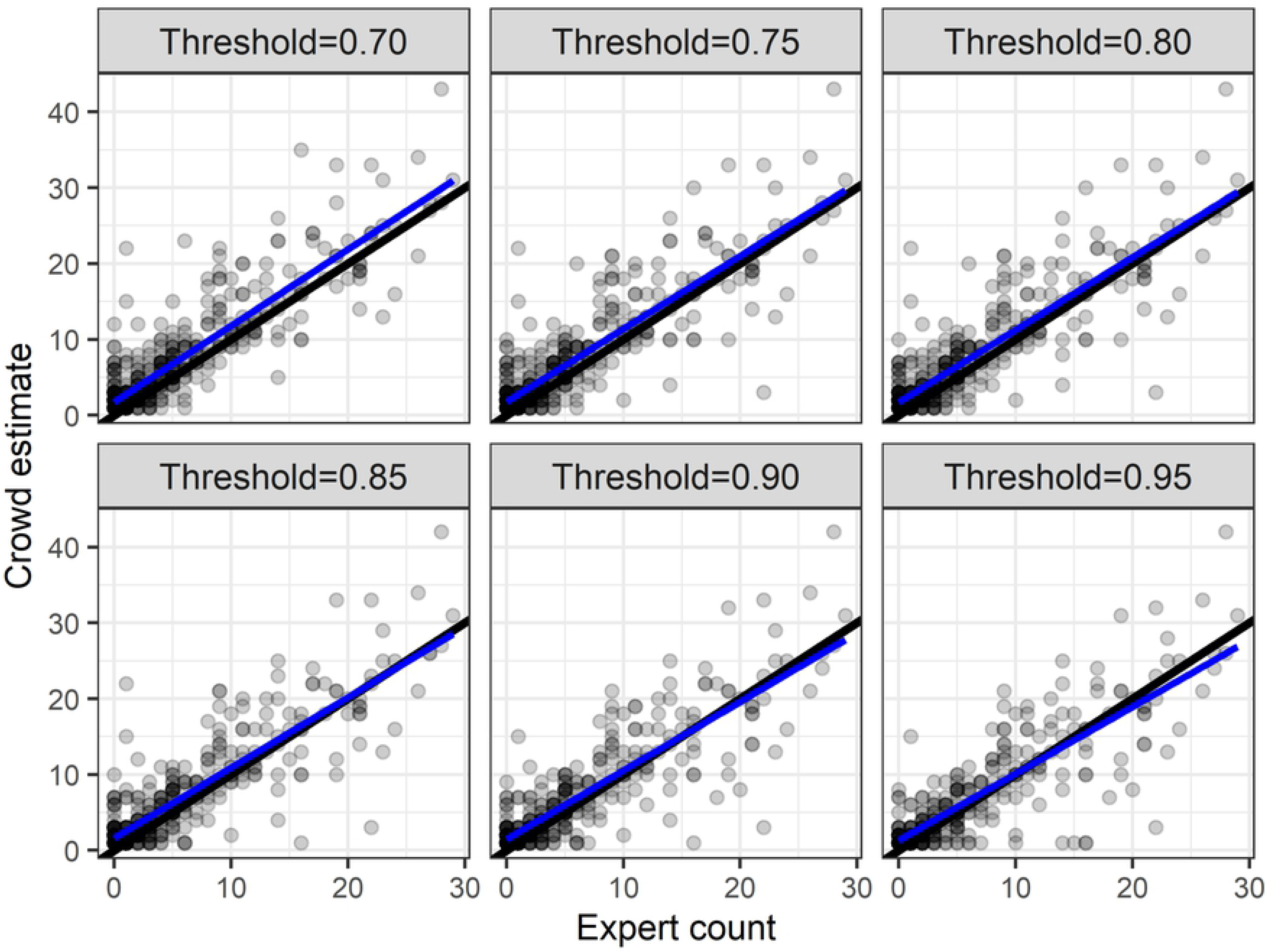
Effect of an increasing threshold filter for feature CrowdRank on the relationship between crowd counts and expert counts. Each point represents the count of an image; darker points indicate overlapping values. The blue line is the best fit linear regression and the black line is the 1:1 relationship line, shown for reference.

Only 4,500 (of 292,049) maps for Erebus Bay had tags from more than one tagger, and so, only these maps contained features. Filtering by feature CrowdRank did not shrink the estimates to values comparable to those of the expert (Fig 2). Visual inspection of Fig 2 shows that increasing the threshold filter for features in a map does not improve the precision or accuracy of the counts. Table 1 compares model fit for different threshold values as filters. We found that the adjusted R-square of the regression model drops with increasing threshold value, contrary to our expectations. Moreover, as the threshold filter increases the number of maps with features assumed to be seals drops. Since the expert tagged 2,854 seals on maps also inspected by the crowd without applying any filters, the use of a feature CrowdRank filter results in many maps with seals having a count of 0 and, for those maps with features above the threshold, counts significantly higher than the expert estimates. Hence, we deemed the “crowd wisdom” method for estimating the number of seals in a map to be unreliable.

**Table 1.** Relationship between the adjusted R-square of the regression of number of features and number of seals in maps, and the threshold filter for feature CrowdRanks.

| Threshold filter | Adjusted R-square | Slope | Intercept | # Maps | Expert count | Crowd estimate | St. Err. Pred. Est. |
| --- | --- | --- | --- | --- | --- | --- | --- |
| 0.7 | 0.78 | 1.01 | 1.73 | 476 | 2355 | 3197 | 96.89 |
| 0.75 | 0.75 | 0.96 | 1.76 | 445 | 2290 | 2986 | 98.46 |
| 0.8 | 0.76 | 0.96 | 1.69 | 440 | 2287 | 2931 | 96.42 |
| 0.85 | 0.75 | 0.93 | 1.61 | 390 | 2158 | 2632 | 94.40 |
| 0.9 | 0.74 | 0.91 | 1.46 | 368 | 2132 | 2472 | 92.73 |
| 0.95 | 0.71 | 0.88 | 1.30 | 326 | 2081 | 2258 | 94.11 |

The use of our expert correction approach to shrink counts resulted in increased precision and accuracy when we filtered data using increasing values of surveyor CrowdRank. Fig 3 shows that precision (as indicated by the size of the error bars around the estimates) and accuracy (proximity of points to the 1:1 line) improve as we filter for higher surveyor CrowdRank. However, filtering comes at the cost of having fewer estimates (because the pool of maps inspected by the surveyors that meet the threshold also decreases). Table 2 shows that as we increase the threshold value, the adjusted R-square of the regression increases and the slope approaches 1. We chose a threshold of 0.8 to strike a compromise between the number of surveyors, number of maps they inspected, and the adjusted R-square of the regression. All results henceforth are obtained after applying the expert correction method obtained from the sample of surveyors meeting (or exceeding) this CrowdRank threshold value. Table 2 shows that this threshold choice results in a representative estimate of the shrinkage factor. On average, the number of seals the expert found in a map is 4.7 times fewer than the features tagged by the crowd of surveyors with surveyor CrowdRank score ≥ 0.8. Even after the shrinkage, and without any further corrections, there is large variance in the estimates from the crowd as compared to the ground counts. We fit the haul-out location and islet/mainland regression models to explore the sources of that variance.

**Fig 3.**
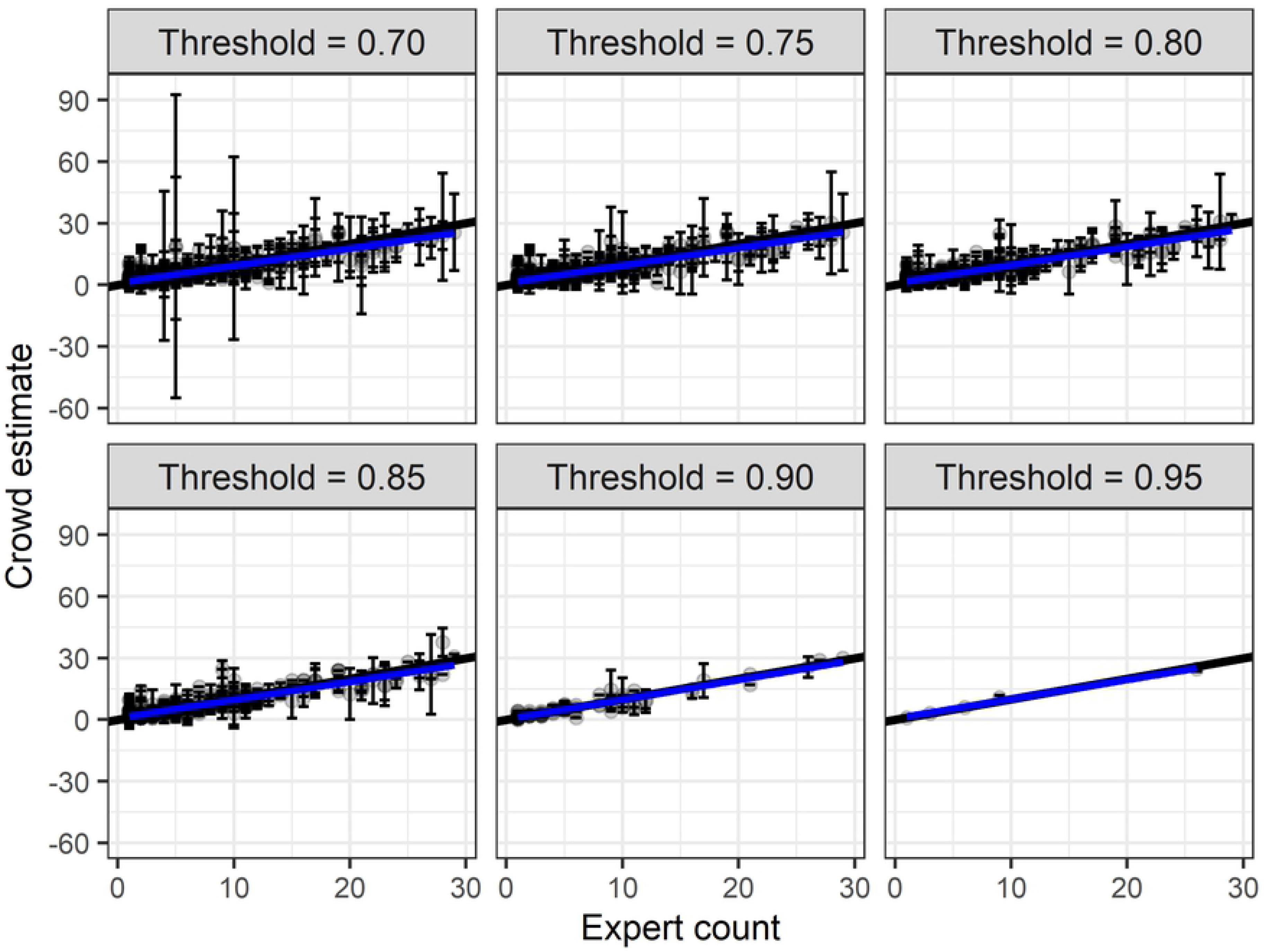
Effect of an increasing threshold filter for tagger CrowdRank on the relationship between bias-corrected crowd counts and expert counts. Each point represents the count of a map; darker points indicate overlapping values. The blue line is the best fit linear regression and the black line is the 1:1 relationship line, shown for reference.

**Table 2.** Effect of an increasing threshold filter for tagger CrowdRank on the adjusted R-square of the regression of bias-corrected crowd counts and expert counts. The crowd estimate is the value corrected by shrinking tagger counts based on their performance against the expert.

| Threshold filter | Adjusted R-square | Slope | Intercept | # Maps | Expert count | Crowd estimate | St. Err. Pred. Est. | Per-image shrinkage |
| --- | --- | --- | --- | --- | --- | --- | --- | --- |
| 0.70 | 0.78 | 0.84 | 0.86 | 443 | 2640 | 2608 | 76.22 | 5.39 |
| 0.75 | 0.82 | 0.86 | 0.84 | 399 | 2440 | 2433 | 67.68 | 4.38 |
| 0.80 | 0.82 | 0.89 | 0.75 | 319 | 2003 | 2023 | 62.75 | 4.70 |
| 0.85 | 0.82 | 0.89 | 0.77 | 306 | 1963 | 1980 | 63.65 | 4.58 |
| 0.90 | 0.91 | 0.96 | 0.15 | 70 | 451 | 445 | 22.28 | 4.90 |
| 0.95 | 0.98 | 0.92 | 1.17 | 3 | 38 | 38 | 3.81 | 1.00 |

The final regression model with haul-out location as a factor is shown in Table 3. The final islet/mainland model is shown in Table 4. The haul-out location model had an adjusted R-squared value of 0.65 (goodness-of-fit data shown in the Supporting Information S1 document). There were no apparent temporal (early versus late in November) effect. Several parameters showed strong effects: location, number of tags placed, number of maps with tags (and the interaction of number of maps with number of tags), and the quadratic of the sinusoidal of hour (with modulus 12 hours) when the image was taken. Both number of tags and number of maps with tags had strong positive effects, thus providing evidence that the detection rate per seal (as determined by ground counts) is higher when the number of tags placed is higher and the area where these were placed is larger, possibly due to a larger number of seals present in a map as well as sample size (number of maps).

**Table 3.** Final linear model of the ratio of corrected crowd counts/ground counts with haul-out location effects. The following covariates were included in the model: number of tags placed and number of maps inspected by the crowd, sinusoidal hour of the day (sinHour), year as factor, and haul-out location. The base value for Location was Big Razorback, and the base value for year was 2010. Adjusted R-squared: 0.653, F-statistic: 18.94 on 13 and 111 d.f., p-value < 0.0001.

| Coefficient | Estimate | Std. Error | t value | Pr(>\|t\|) |
| --- | --- | --- | --- | --- |
| (Intercept) | 0.246 | 0.025 | 10.018 | < 0.001 |
| Number of tags | 0.086 | 0.009 | 9.390 | < 0.001 |
| Number of maps | 0.038 | 0.010 | 3.793 | < 0.001 |
| Number of tags x number of maps | -0.023 | 0.006 | -3.884 | < 0.001 |
| sinHour | -0.008 | 0.012 | -0.656 | 0.513 |
| sinHour^2 | -0.040 | 0.019 | -2.138 | 0.035 |
| Year (2011) | -0.061 | 0.022 | -2.790 | 0.006 |
| Location: Hutton Cliffs | -0.124 | 0.023 | -5.463 | < 0.001 |
| Location: Inaccessible Island | 0.122 | 0.024 | 5.028 | < 0.001 |
| Location: Little Razorback | 0.221 | 0.024 | 9.145 | < 0.001 |
| Location: South Base | 0.088 | 0.048 | 1.843 | 0.068 |
| Location: Tent Island | -0.032 | 0.021 | -1.550 | 0.124 |
| Location: Turks Head-Tryggve | -0.089 | 0.022 | -3.952 | < 0.001 |
| Location: Turtle Rock | 0.112 | 0.020 | 5.614 | < 0.001 |

**Table 4.** Final linear model of the ratio of corrected crowd counts/ground counts with islet/mainland effects. The following covariates were included in the model: number of tags placed by the crowd, sinusoidal hour of the day (sinHour), year, and islet/mainland effects. Adjusted R-squared: 0.306, F-statistic: 11.91 on 5 and 119 d.f., p-value < 0.0001.

| Coefficient | Estimate | Std. Error | t value | Pr(>\|t\|) |
| --- | --- | --- | --- | --- |
| (Intercept) | 0.152 | 0.034 | 4.538 | < 0.001 |
| Number of tags | 0.062 | 0.009 | 6.604 | < 0.001 |
| sinHour | -0.016 | 0.017 | -0.936 | 0.351 |
| sinHour^2 | -0.045 | 0.026 | -1.714 | 0.089 |
| Year (2011) | -0.044 | 0.030 | -1.502 | 0.136 |
| Islet | 0.115 | 0.019 | 5.929 | < 0.001 |

Likelihood ratio tests (Likelihood Ratio Statistic = 125.39, 7 d.f., p < 0.001) indicated that location (i.e., of the haul-out aggregation) accounted for approximately half of the variance explained. Table 3 also shows that, contrasted with Big Razorback (the reference site in the model, an “islet” location), the “mainland” sites, Hutton Cliffs and Turks Head-Tryggve, showed very strong negative effects on detection rate. That is, detection rate estimates from these “mainland” sites (i.e., near Ross Island in these two cases) are much lower than those of other locations (Table 3), supporting the dichotomy explored in the islet/mainland model.

Table 4 shows the results of the final islet/mainland model. Under this model, there is no longer an effect of number of maps with tags, or of its interaction with number of tags placed. This indicates a confounding effect of number of maps and the islet/mainland binary. Indeed, the mainland locations had larger numbers of seals. The model with islet/mainland effects had an adjusted R-squared value of 0.31, which was less than half that of the haulout-location model (Table 3) (see also goodness-of-fit plots shown in the Supporting Information S1 document).

Fig 4 shows the predicted abundances for each aggregation using the above two models (A: haul-out location model; B: islet/mainland model). The diagonal line represents the best fit from the regression model between the estimate and the ground count. There is overall agreement in estimates for the smaller islet locations with both approaches but not so for one of the larger mainland locations in one of the models. The estimated value for Turks Head-Tryggve on one survey occasion using the islet/mainland model was quite misleading: we calculated <50 seals when in fact there were three times that many on that date (as determined by ground counts).

**Fig 4.**
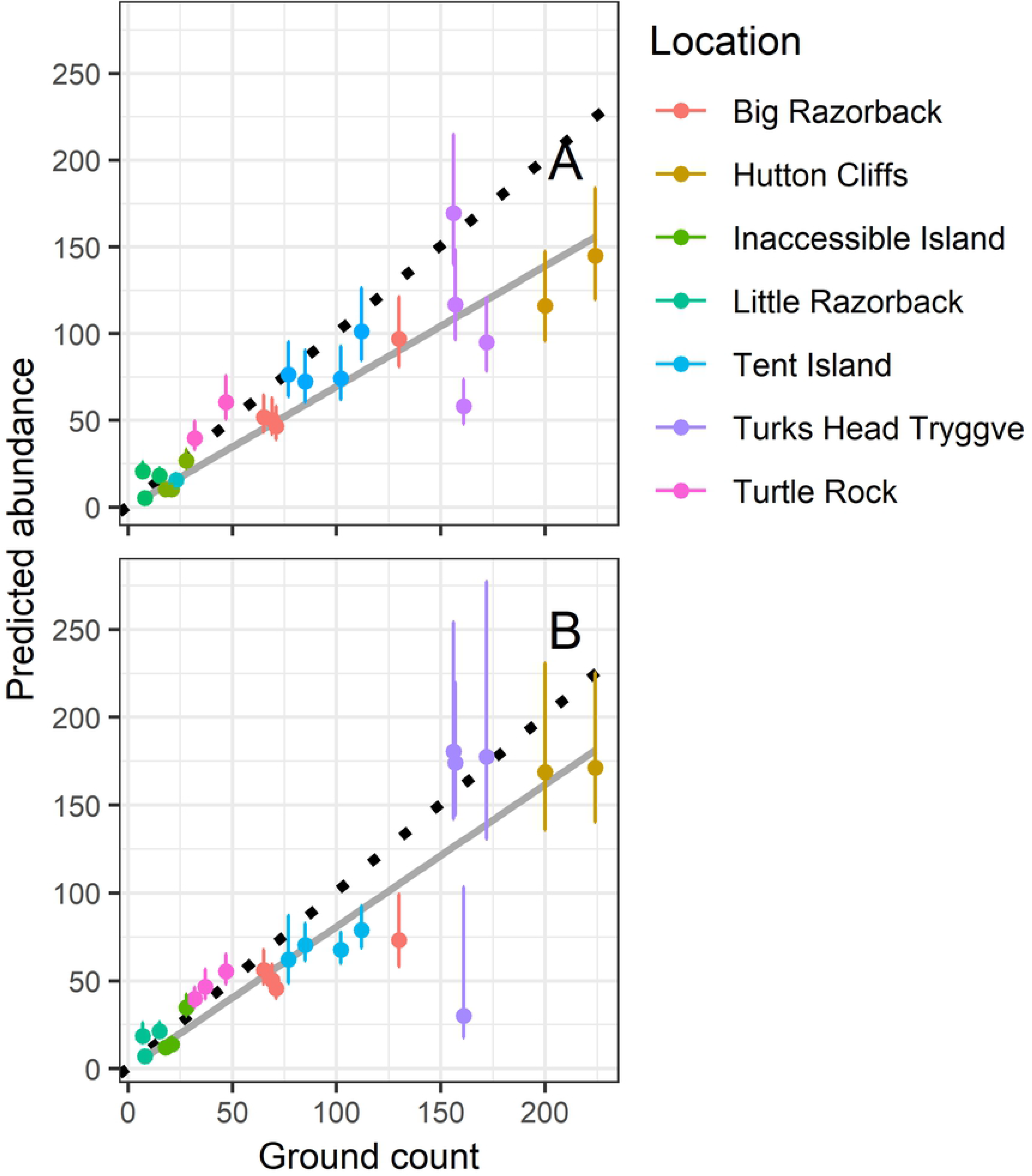
Estimated number of seals using the predicted detection rate from the best model that accounts for either (A) haul-out location effects (8 locations) or (B) islet/mainland effects, as compared to the ground counts. The bars are the inner 95-percentile values of the estimates. The solid gray line is the fitted linear model of estimated abundance vs ground counts. The 1:1 relationship line (dotted) is shown for reference.

By scaling up to all of Erebus Bay, we found that predictions of abundance from the islet/mainland model performed well. Fig 5 shows that model-predicted abundance for Erebus Bay (solid line) closely approximated the 1:1 line (dotted line) and that most of the variation in abundance for Erebus Bay for each satellite image was accounted for by variation in model-predicted abundance. Since satellite images never encompassed all haul-out locations, the predicted abundances vary due to both the normal variation in aggregation numbers over time, as well as the specific combination of haul-out locations included in each satellite image.

**Fig 5.**
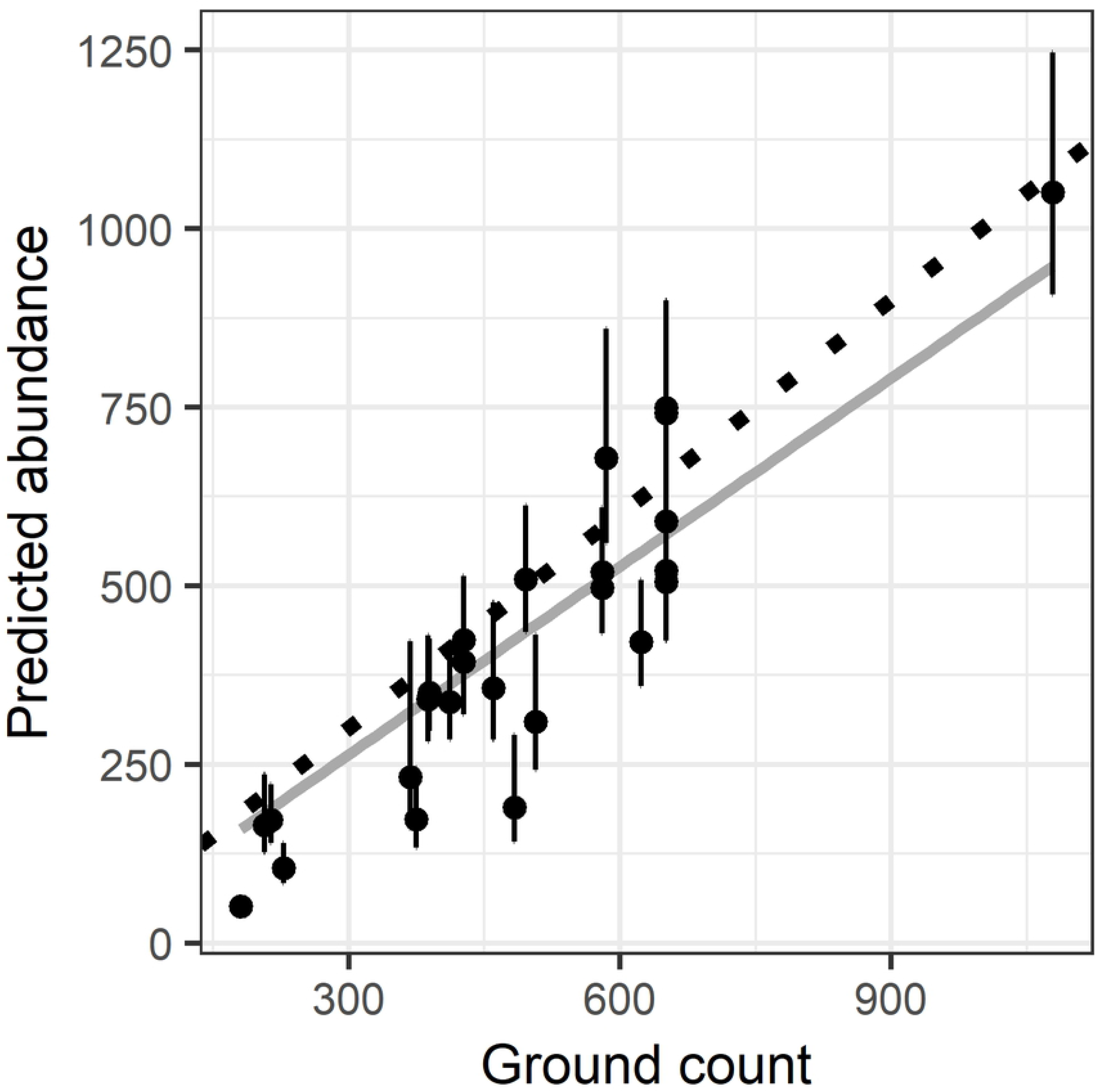
Estimated number of seals from the islet/mainland model at the regional Erebus Bay scale. Each point is the observed and predicted abundance of seals from all locations in each photograph combined. The solid gray line is the fitted linear model of estimated abundance vs ground counts. The 1:1 relationship line (dotted) is shown for reference.

## DISCUSSION

### Understanding the bias in citizen scientist counts

Responding to our request via the Tomnod platform, >240,000 citizen scientists counted Weddell seals in satellite images. Each map was inspected by as few as two (18% of images, the most common case) and as many as 15 citizen scientist taggers (0.04% of images); 88% of the images were inspected by 2-5 citizen scientists. By comparing individual tagger counts against an expert, we were able to detect a counting bias by the crowd (i.e., higher counts by taggers than found by the expert), as well as variance among taggers. We tested two alternative ways to estimate and correct for the bias. Below we review our choices of methods, as well as other possible solutions.

Our study design followed the approach described by Kamar et al. [46] to understand and address data quality. Using the Tomnod algorithm, we estimated the tagger CrowdRank value and simultaneously the feature CrowdRank value for each feature tagged by at least two citizen scientists. We used the feature CrowdRank to decide if a map should remain in the pool for further examination, and the tagger CrowdRank to measure the degree to which other taggers agree with his/her determination that a feature is a seal. However, the tagger CrowdRank is just an index of agreement among taggers. It is not an index of the correct determination. As such, it operates on an underlying rate of misidentification of features as seals common to all taggers, so that often features other than seals end up with very high CrowdRank values. In other words, it is not enough to use tagger agreement to determine if a feature is a seal or not. Our results show that on numerous occasions, features on a map other than seals end up being highly ranked, or conversely, that true seals were not counted. Therefore, this approach did not provide reliable means for the crowd to self-correct and reduce sampling bias. This bias in surveyor skills is a drawback not exclusive to citizen science research [16], but it can be a limiting factor of citizen science if not properly addressed [16,47]. With our approach, external information was needed to properly correct the error introduced by the citizen scientists.

We show that the calibration against an expert to develop a correction factor (involving a “shrinkage” factor) is able to account for the imperfect detection and probability of false positive assignments among surveyors (see the Supporting Information S1 document for details). This approach provides reasonably accurate and unbiased estimates of seals in maps. However, the number of seals detected in maps, even by the expert, is only about 2/3 of all seals actually on the ground (Fig 4). This difference is not necessarily due to taggers (or the expert) failing to detect seals in an image, but more likely due to the fact that some fraction of seals will be in the water, and not hauled out on the ice [48,49]. The proportion of seals diving versus on the ice varies throughout the day, but with a discernible (and thus correctable) pattern ([48,49]; this study).

### Other possible approaches to reducing bias

A more direct solution to the sampling bias problem would be fitting hierarchical models with a latent count variable to estimate a probability of imperfect detection (i.e., not counting seals present in an image), and a simultaneous probability of a false positive determination of a seal from a feature. This approach has been used for the estimation of species presence (e.g., [50]), following a model first proposed by Royle and Link [51]. Since the false negative and false positive probabilities operate simultaneously on the detection process, estimating them from data requires a model parametrization that includes covariates informing the seal detection process, and covariates informing the assignment of false positives. The covariates of detection and of false positive assignments would be such that they would vary among taggers. Unfortunately, we lack any covariate data to inform a model on differences among taggers, other than their performance among peers and, for a fraction of them, their performance versus the expert.

Our method is unique in that it simply requires a representative selection of participants to be calibrated against an expert to develop a correction factor that is applicable to the entire set of participants. We do not limit the scope of participation based on some prior knowledge of surveyor quality. We have not attempted an approach in which every participant is engaged first in a calibration against an expert, which would then provide us with some *a priori* information on tagger quality that can be used with our methodology or possibly with hierarchical models. Another approach, proposed by See et al. [52], includes the use of training materials and feedback for the purpose of increasing the quality of data from the crowd. When the number of participants is large, the feedback process may be costly. In our case, we included only some very limited and easy-to-follow training materials. More complex training may be a deterrent for participation, but we concur with the authors that better training of participants will readily result in higher quality of data and more precise estimates.

### Comparing counts in images versus ground counts

Available on-the-ground haul-out location counts allowed us to correct for the proportion of seals not detected through satellite images. Still, the mainland aggregations were underestimated (addressed below). Although our “expert-correction” method accounts for the seals’ known daily diving patterns [40,45,49], it does not correct for the difference between calibrated crowd counts and ground counts. This is because there is always a proportion of seals (apparently about 1/3 of them) not on the ice and not available for detection. The hour effect is modeling how this proportion varies during the day. This hour effect was evident in both the haul-out location model and the islet/mainland model. There is also a small year effect in both models (significant only in the haul-out location model, which is the better fit). This effect indicates that the detection rate was slightly lower in 2011 as compared to 2010. The number of tags placed and number of maps inspected per location helped explain some of the variance of the haul-out location model. Larger haul-out locations host more seals and require more maps to inspect by the crowd. Similarly, both models show that detection rate correlates with total number of tags placed per map, as evidenced in a positive slope for total number of tags placed.

Our ultimate objective is to estimate the number of seals throughout Antarctica, but since we have only estimates for haul-out aggregations in Erebus Bay, we cannot impute specific haul-out location effects from aggregations in Erebus Bay to other unknown locations. Moreover, we lack any *a priori* knowledge of aggregations in other locations to apply location effects from a “matching” aggregation in Erebus Bay. This is why, instead, we used a characteristic that could be used throughout Antarctica: the islet/mainland effect. Table 3 indicates that locations within Erebus Bay differ substantially with respect to detection rate, even when comparing among mainland or among islet sites. Without taking into account specific location, and instead modeling an islet/mainland effect, we obtained a useful predictive model but with reduced explanatory power. The adjusted R-square of the haul-out location model is 65%, versus 31% of the islet/mainland model. Fig 4B shows the predicted values using the islet/mainland model directly, which, with the exception of one outlier, performed relatively well. Ideally, we would like to test this approach with islet and mainland aggregations outside of Erebus Bay.

Fig 5 shows that the loss of explanatory power can be reduced if counting seals at scales larger than the individual haul-out aggregation. This approach thus sacrifices spatial specificity for accuracy at the larger scale, but is a promising means for surveying the entire Weddell seal populations for all of Antarctica, a subject which we address next.

### Monitoring Weddell seal populations for the entire continent

Does our methodology provide us with a tool to monitor seal populations locally, regionally, or continent-wide? Since Erebus Bay uniquely hosts the largest accumulation of seals anywhere in Antarctica (our unpublished results), does this invalidate the application of our methods to other regions of Antarctica? It is possible that the error committed by the crowd increases proportionately with the size of the local aggregation of seals. If that is the case, it is possible that our approach cannot simply be extended to the rest of the continent if clusters of seals elsewhere are much smaller than those found in Erebus Bay. Based on preliminary results, we have no basis to suspect that this is the case.

Other sea-ice or polar species have been studied through satellite images. Fretwell et al. [53] showed that satellite imagery can be used to estimate the global population of the Emperor penguins (*Aptenodytes forsteri*) in Antarctica. Using the same methodology, Lynch and LaRue [54] estimated the global population of Adélie penguins (*Pygoscelis adeliae*) in Antarctica, and Stapleton et al. [55] estimated polar bear (*Ursus maritimus*) numbers in the Arctic. These other studies were all conducted by researchers, not the crowd. But some research problems may require large teams or crowd sourcing if the data stream that is to be analyzed is quite large (as in our case). Indeed, crowdsourcing and citizen science can be the next step in a progression from individual researchers, to large research teams, to large crowds, enabling scientists with the capacity to answer questions unattainable under other research models [56]. Our research goal fall squarely in this last category.

Our results indicate that it is indeed feasible to monitor Weddell seal populations at least regionally, if not continent-wide, using our methodology and citizen science. Indeed, our study provides evidence that citizen science can contribute to a rigorous scientific endeavor on par with conventional scientific methods, and effectively inform conservation action [47]. And if the count estimates from our methods are imprecise at a particular spatial scale, some degree of monitoring would still be informative if we develop indices of relative abundance.

In this study we used 2 years of very high resolution satellite imagery to test and validate our methodology (e.g., Fig 7). There are many more years of imagery available, thus employing the crowd to assess a longer time series appears feasible. Such an approach would be needed for example to adequately monitor seal populations within the context of conservation and impact assessment of long-term drivers, such as a fishery or climate change. CCAMLR is mandated to monitor “related or dependent species” relative to any extraction activities, thus, to minimize or avoid adverse impacts. For example, the CCAMLR Ecosystem Monitoring Program (CEMP) was designed to monitor trends in land-based species that could be affected by a fishery for Antarctic krill (*Euphausia superba*) [57,58]. But the organization has yet to extend the effort to monitoring effects of finfish fisheries. If the toothfish fishery is having an impact on Weddell seal populations, a real possibility given that toothfish contribute significantly to the seals’ diet [25,59], we may be able to discover a correspondence between seal abundances at various locations in the Ross Sea and extraction numbers or some metric of fishing effort [33]. We could then contrast these patterns with other regions or time periods where the fishery is not (yet) operating to better inform conservation and regulation. The methodology presented here, given adequate funding and a citizen science platform like Tomnod, would indeed allow us to do just that, monitor Weddell seals from space to better inform conservation and regulation.

### Correcting citizen science-based wildlife counts

Our “shrinkage factor” calculation can be applied to a broad range of wildlife count data. Any citizen science count dataset is likely to be subject to variance in surveyor skill and the potentially biased distribution of these skills in place and time. As long as an expert assessment is available and can be contrasted to a representative subset of the surveyors to determine the accuracy of their counts, our method can be used to remove the bias and correct the counts. We show that this approach produces estimates that can be sufficiently accurate at the appropriate spatial scales. We also show that it is a much more reliable method than those that rely on how surveyors rank each other (“crow wisdom”). There is nothing intrinsic to the data that guarantees it is free of bias in surveyor skill. Careful examination and estimation of the bias to correct it must be a fundamental component of citizen science-based wildlife counts. The method presented here is a step in that direction.

## ACKNOWLEDGMENTS

We greatly appreciate the help from thousands of volunteers who participated in this project. We wish to thank: the Antarctic and Southern Ocean Coalition; the University of Minnesota, DigitalGlobe Foundation, and SciStarter for their support; the Polar Geospatial Center for assistance in gathering image data; and the graduate students and field technicians who collected ground survey data. Logistical support for fieldwork was provided by Leidos, Lockheed Martin, Raytheon Polar Services Company, Antarctic Support Associates, the United States Navy and Air Force and Petroleum Helicopters Incorporated.

